# Zebrafish prph2a/b and rom1a/b serve distinct functions during cone and rod outer-segment assembly

**DOI:** 10.64898/2026.08.06.743304

**Authors:** Meet Patel, J.K Famulski

## Abstract

Inherited retinal disorders are significant contributors of blindness worldwide. Mutations in Peripherin-2 (PRPH2), a highly conserved vertebrate tetraspanin membrane protein responsible for formation and maintenance of OS morphology, have been shown to cause diverse types of inherited photoreceptor cell (PRC) disorders including but not limited to Leber congenital amaurosis, cone-rod dystrophy, and retinitis pigmentosa. In this study we used a cone-rich diurnal zebrafish model to characterize the loss of PRPH2 function. Of the four PRPH2 zebrafish orthologs only prph2a and prph2b were found to be expressed in PRCs. CRISPR-mediated single mutants of prph2a and prph2b did not yield striking rod or cone phenotypes. Double prph2a/2b mutants exhibited early loss of all cone cells, preceded by cone outer segment disorganization in the form of whorls akin to the phenotypes observed in PRPH2+/- mice. Surprisingly rod photoreceptor cells were not affected and in fact exhibited a striking lengthening of rod OSs with normal disc formation. Overgrowth of rod OSs proceeded up to 1 year, but no degeneration was observed. To determine how rod OS can persist without prhp2a/b we targeted rom1a and rom1b using CRISPR. Injection of rom1a/b crRNA resulted in complete loss of both rod and cone OSs in the prph2a/b double mutants. Surprisingly, inhibition of rom1a/b alone resulted in the loss of rod but not cone OSs. These findings suggest that unlike in mammals, zebrafish rom1a/b is essential for rod OS formation while prph2a/b is essential for cone OSs.

## INTRODUCTION

Vertebrate retina consists of three cell layers including the outer nuclear layer which contains rod and cone photoreceptors, and functions to convert light into electrical signals so visual stimuli can be subsequently detected by various cortical regions of the brain^1^. Phototransduction is carried out in the Outer Segment (OS) of both rod and cone PRCs. In rods the mature OS is made up of discrete orderly stacked discs like thylakoids in chloroplasts encased by the cell membrane whereas immature rod OS discs and majority of the cone OS discs are open to the extracellular matrix and are continuous with the cell membrane^2^. Both rod and cone OS discs contain their respective opsin and voltage gated ion channels required for phototransduction^3,4^. Due to constant bombardment of photons, the OS remains a highly metabolic environment, and thus OS discs are renewed periodically as a protective mechanism to avoid oxidative stress^5,6^. Old OS discs are constantly phagocytized and degraded apically by the Retinal pigmented epithelium (RPE) which demands a constant supply of new OS discs distally ^5,7^. In rod OS, discs are first produced as evaginations from the connecting cilium at the base of the OS and require PRPH2, to localize at the outer rims of rod OS discs to drive disc rim biogenesis via formation of PRPH2 oligomers. PRPH2 function is essential to transform these evaginations into an ordered pattern of OS discs^8–10^. Although the evagination process in formation of cone OS is largely thought to be consistent with rod OS, the localization of PRPH2 and thereby its role appears to be different in cone OS biogenesis compared to rods^9^. As such the complete function of PRPH2 in cone OS biogenesis and homeostasis is not thoroughly understood.

Roughly 5% of all the diagnosed Inherited Retinal diseases are PRPH2 associated retinal diseases (PARDs) resulting from pathogenic PRPH2 variants^11,12^. These mutations primarily affect the Photoreceptor and the RPE cells^11^. Numerous studies have highlighted the phenotypic variability due to perturbations in PRPH2 function leading to IRDs including but not limited to Retinitis Pigmentosa, macular dystrophy, Leber Congenital Amaurosis (LCA), and cone rod dystrophy^12–15^. As such, the burden of PARDs bears the need to elucidate the differential role of PRPH2 in the rod and cone photoreceptor cells as well as the indirect effects on RPE.

Structurally, PRPH2 consists of four helical transmembrane domains, three cytoplasmic domains, and two intradiscal (in the OS disc lumen) domains^8^. Via the larger 2^nd^ intra discal domain, it binds with other PRPH2 to result in PRPH2 oligomers^8,9^. In cone OS these PRPH2 oligomers generate disc rim curvature specifically at the edge closest to the connecting cilium where its localization is restricted^9,16^. Based on localization studies, it is speculated that PROM1 in cone OS may be responsible for performing a similar role as PRPH2 but on the opposing side of the connecting cilium mirroring PRPH2 localization^17,18^. In rod OS, although PRPH2 makes PRPH2 based oligomers, it also interacts with a PRPH2 homolog tetraspanin ROM1 by forming heteromeric oligomers via interactions with their respective 2^nd^ intra discal domains resulting in rod OS rim curvature^19,20^.

Loss of a single functional copy of PRPH2 in heterozygous mice leads to formation of OS whorls while mice with complete loss of PRPH2 fail to form any OS disc in rod or cone, and instead only exhibit OS vesicles^21–24^. Further, several mutant variant mice models have shown residue level resolution of PRPH2 function with majority of the key residues predominantly involved in higher order oligomer formation residing in the 2^nd^ intradiscal domain while some key residues in the C terminus responsible for its non-canonical trafficking trail to the OS^25–29^. These mice models have been vital in our understanding of PRPH2 function. However, since majority of PRPH2 studies have employed the mouse model, which is rod dominant due to its nocturnal nature, there is a significant lack of efforts to examine PRPH2 homologs in diurnal models as to better understand cone PRC pathophysiology and the spectrum of phenotypes seen in the clinic. As such, we sought to develop a cone-rich diurnal zebrafish model to characterize the loss of PRPH2 function and compare to the diurnal mouse model.

In the current study we determine that in zebrafish two PRPH2 orthologues contribute to OS function, prph2a and 2b. We show that inactivation of both prph2a and 2b results in the loss of cone, but not rod, OS integrity leading to degeneration. Furthermore, we show rod specific compensation of prph2 function by rom1a and rom1b and the sole necessity of rom1a and 1b during rod OS formation.

## RESULTS

### Prph2a and prph2b are zebrafish orthologs of PRPH2

PRPH2 deficiency has been extensively modelled using the murine model which recapitulates several human phenotypes including retinitis pigmentosa and choroidal dystrophy^21,30^. However, PRPH2 deficiency has yet to be studied in a diurnal or a cone rich model. As such, we sought to model PARD using the cone-rich diurnal zebrafish. The zebrafish genome encodes four prph2 orthologs: *prph2a, prph2b, prph2la*, and *prph2lb* respectively. AlphaFold structure predictions of these orthologs suggests a strikingly similar molecular structure despite only ∼60% residue conservancy (Fig S1A). The predicted structures possess the same PRPH2 homologous structures including four transmembrane domains, cytoplasmic N and C termini and cytoplasmic loops.

Further, all the orthologs encode two intradiscal domains including the critical (D2) 2^nd^ intra discal domain. As such all four orthologs could potentially to contribute to prph2 function in zebrafish PRCs. However, using Whole Mount In situ hybridization (WISH), we only observed the expression of *prph2a* and *prph2b* in photoreceptors at 5dpf (Fig S1B). Expression of *prph2la* was absent from the retina while we were unable to detect any *prph2lb* mRNA at 5dpf or older stages (Fig S1B). Additionally, when examining the daniocell database (https://daniocell.nichd.nih.gov/), and an adult single cell RNA data set we confirmed *prph2a* and *prph2b* expression in both cone and rod cells ^31^. As such, we conclude that *prph2a* and *prph2b* are the only retinal prph2 zebrafish orthologs. Thus, we next sought to evaluate the phenotypes associated with the loss of either prph2a or prph2b in the cone-rich diurnal zebrafish retina.

### Individual prph2a or prph2b loss of function mutations have minor effects on zebrafish photoreceptor OSs

Since prph2a and prph2b are the only PRPH2 orthologs expressed in the zebrafish retina, we generated loss of function alleles in each using Alt-R CRISPR technology. For prph2a, we used two crRNA constructs separated by 274bp and mapping to exon 1. Upon injection, we generated a stable prph2a mutant line harboring a 359bp deletion resulting in a frame shift at AA69 followed by a premature stop codon at AA90 (Fig 1A). Wildtype prph2a protein sequence consists of four transmembrane, three cytoplasmic and two intradiscal domains, while the mutant transcript results in a truncated polypeptide consisting of the N terminus, the first two transmembrane domains, the first intradiscal domain, and a malformed initial cytoplasmic domain truncating at the AA90. Thus, any remaining protein would lack both the critical 2^nd^ intradiscal domain and the signaling C terminus domain. The prph2a^fs69–90*^ line (referred to as prph2a^-/-^) was bred to homozygosity to create a maternal zygotic prph2a^-/-^ mutant line. These maternal zygotes were used for all the subsequent experiments. Using WISH, we determined that the prph2a mutant allele mRNA undergoes non-sense mediated decay further validating it as a null allele (Fig S2A-A”).

**Figure 1:**
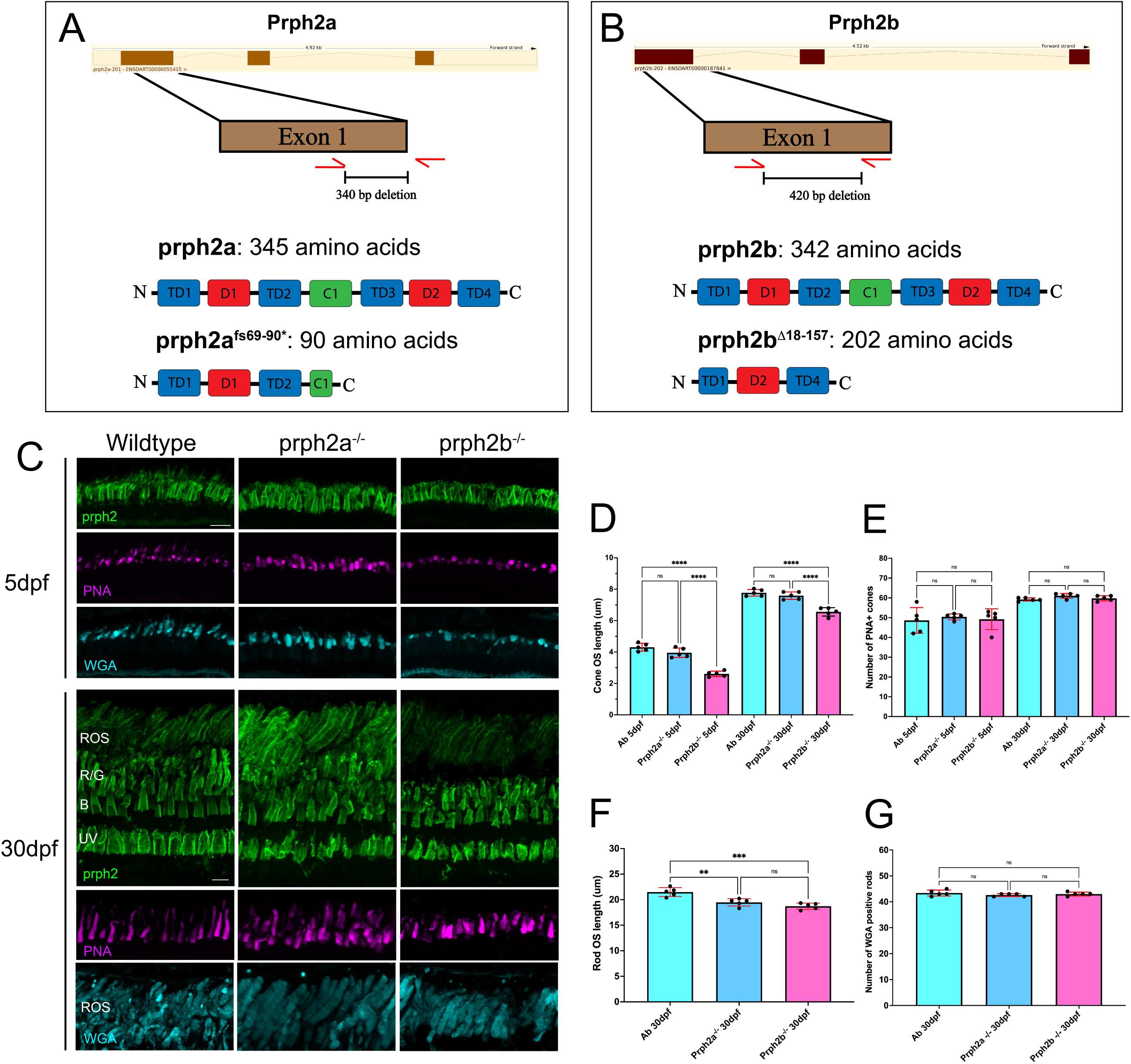
Single prph2a and prph2b zebrafish mutants display minor photoreceptor outer segment phenotypes. **A)** Schematic outlining the CRISPR-Cas9 generated prph2a mutant line harboring a 340bp deletion resulting in a frame shift at amino acid 69 and a premature stop codon at amino acid 90. **B)** Schematic outlining the CRISPR-Cas9 generated prpb2b mutant line harboring a 420bp deletion resulting in an in-frame deletion of amino acids 18-157. **C)** Confocal imaging of central retina cryosections from 5 or 30dpf wildtype prph2a-/- or prph2b-/- individuals, probed with anti-prph2 antibody (green), PNA to detect green cones (magenta) and WGA to detect red cones and rods (teal). ROS = rod outer segments, R/G = red/green cones, B = blue cones, UV = UV cones. Scale bar = 5μm. **D)** Quantification of cone OS length. * p<0.05, ** p<0.01, *** p<0.001, **** p<0.0001. **E)** Quantification of cone cell number at 5 and 30dpf. **F)** Quantification of rod OS length at 30dpf. **G)** Quantification of rod cell number at 5 and 30dpf. * p<0.05, ** p<0.01, *** p<0.001, **** p<0.0001.

The prph2b loss of function mutant line was similarly generated using the Alt-R CRISPSR strategy. Here the two crRNA constructs were separated by 434bp also mapping to exon 1. Post injection, we established a stable prph2b mutant line harboring a 420bp deletion resulting in an in-fame deletion of AA18–157 (Fig 1B). Consistent with prph2a, the wildtype prph2b polypeptide consists of four transmembrane, three cytoplasmic and two intradiscal domains, while the mutant prph2b transcript leads to a shortened polypeptide consisting of the N terminus, a truncated 2^nd^ intradiscal domain, a single terminal transmembrane domain, and the C terminus. We therefore predict that the deletion results in a non-functional truncated prph2b polypeptide that is lacking the first three transmembrane domains and the 2^nd^ cytoplasmic loop. We refer to the allele prph2b^Δ18–157^ as prph2b^-/-^. Unlike prph2a, WISH revealed a persistent expression of prph2b mRNA in the prph2b^-/-^ line indicating a lack of non-sense mediated decay (Fig S2B-B”). Using immunohistochemistry, in both prph2a^-/-^ or prph2b^-/-^ retinas we still detected prph2 signal in the retina, indicating that our commercial antibody likely recognizes both orthologs (Fig 1C). Sequence similarity between prph2a and prph2b is very high so cross-reactivity was not unexpected.

To examine the impact of single prph2a^-/-^ and prph2b^-/^- mutants, we collected retinas at 5 and 30dpf. At each timepoint we examined OS morphology and number of cones and rods cells by labeling the outer segment using prph2 antibodies, Peanut Germ Agglutinin (PNA) and Wheat Germ Agglutinin (WGA) to label cones and rods respectively. We analyzed five individuals of each genotype at each timepoint. All images were captured from the central retina. At 5dpf we observed that WT, prph2a^-/-^and prph2b^-/-^ cone OSs exhibited their expected shape and contained similar numbers of cells (Fig 1D). Upon closer examination we found the average cone OS length measured via PNA staining showed a slight but significant reduction in cone OS length in prph2b^-/-^ at 2.6μm compared to 4.3μm in WT embryos (Fig 1E, S2C). By 30dpf, we continued to detect a reduction in PNA+ cone OS length in prph2b^-/-^ at 6.6μm compared to 7.8μm in WT. There was no effect on the number of cone cells at either timepoint indicating the slight reduction in OS length does not lead to degeneration (Fig 1E).

Taken together, prph2a loss of function had no effect on cones while prph2b loss of function lead to minor decrease in cone OS length but did not impact cone survival. When analyzing rod OSs we again observed no gross morphological phenotypes, but OS length measurements at 30dpf showed slight, but significant decrease in rod OS length for both prph2a^-/-^ and prph2b^-/-^, 19.5μm and 18.7μm vs 21.5μm in wildtype (Fig 1F, S2D). Like with cones, despite the slight decrease in OS length, rod cell numbers in either of the single mutants did not differ from wildtype indicating the OS lenght decrease does not lead to rod degeneration (Fig 1G). The absence of expected severe phenotypes in rods and cones based on mouse studies is not surprising since high conservation between the two zebrafish prph2 orthologs is likely to enable them to compensate for one another. The slight, but significant, decrease in cone OS length in prph2b^-/-^ retinas suggests that the two orthologs may have evolved distinct roles in cone OS formation, with prph2b possibly having a bigger impact on OS length. Rod OS length was affected similarly by the loss of either prph2 ortholog indicating that zebrafish rod OS length may be sensitive to the overall levels of prph2a/b protein rather than a specific ortholog.

### Loss of both prph2a and prph2b leads to cone OS whorls and degeneration

Having observed only minor phenotypes in our single mutant lines, we next established a homozygous prph2a/2b^-/-^ double mutant line. To confirm we have generated a total prph2 knockout we probed double mutant retinas with prph2 antibody. To our surprise, at 5dpf we still detected prph2 signal, however the signal was restricted exclusively to rod cells and completely absent from cone cells (Fig 2A). All prph2 positive cells were also positive for rhodopsin as indicated by the 1D1 antibody while PNA, UV opsin or blue opsin positive cone cells all lacked all prph2 signal (Fig 2B). Based on these results we hypothesized whether the prph2 antibody we used could cross-reacts with the rod specific rom1, a close relative of prph2, which is only expressed in zebrafish rods.

**Figure 2:**
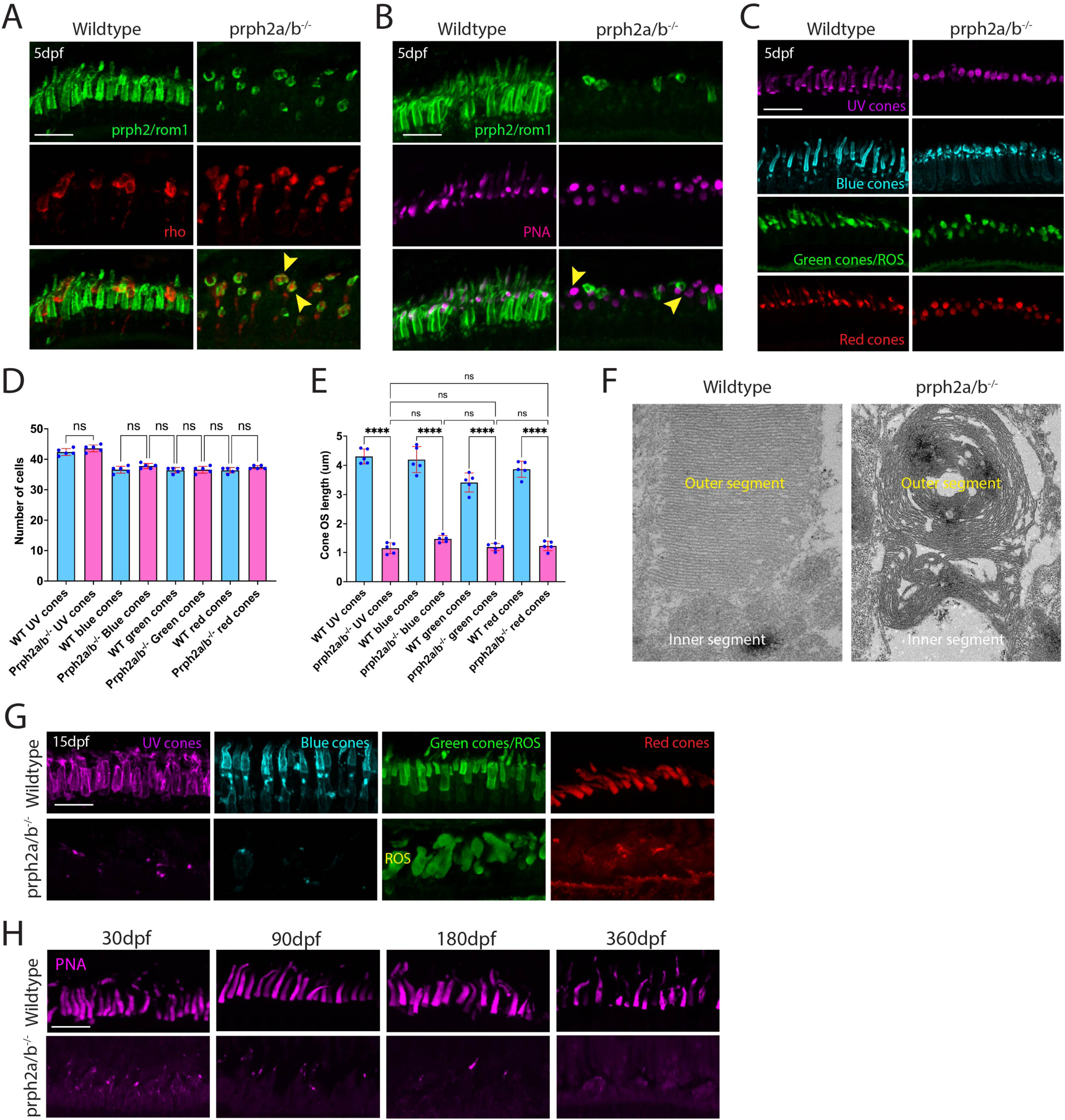
Loss of prph2a and prph2b leads to cone OS whorls and subsequent cone cell degeneration. **A)** Confocal imaging of central retina cryosections from 5dpf wildtype or prph2a/b^-/-^double mutant individuals, probed with anti-prph2 antibody (green) and anti-rho antibody (red). Co-labeling in individual rod cells is highlighted by yellow arrows. Scale bar = 5μm. **B)** Confocal imaging of central retina cryosections from 5dpf wildtype or prph2a/b^-/-^ double mutant individuals, probed with anti-prph2 antibody (green) and PNA (magenta). Absence of co-labeling in individual cone cells is highlighted by yellow arrows. Scale bar = 5μm. **C)** Confocal imaging of central retina cryosections from 5dpf wildtype or prph2a/b^-/-^ double mutant individuals, probed with anti-UV opsin antibody (magenta), anti-blue opsin antibody (teal), WGA (green) or PNA (red). Scale bar = 5μm. **D)** Quantification of the number of each cone cell type at 5 dpf compared to wildtype. * p<0.05, ** p<0.01, *** p<0.001, **** p<0.0001. **E)** Quantification of the OS length of each cone cell type at 5 dpf compared to wildtype. * p<0.05, ** p<0.01, *** p<0.001, **** p<0.0001. **F)** Transmission electron microscopy images of wildtype and prph2a/b-/-retinal sections at 5 dpf. **G)** Confocal imaging of central retina cryosections from 15dpf wildtype or prph2a/b^-/-^ double mutant individuals, probed with anti-UV opsin antibody (magenta), anti-blue opsin antibody (teal), WGA (green) or PNA (red) and anti-rho antibody (red). Scale bar = 5μm. **H)** Confocal imaging of central retina cryosections from 30, 90, 180 and 360dpf wildtype or prph2a/b^-/-^ double mutant individuals, probed with PNA (magenta). Scale bar = 5μm.

Zebrafish prph2 orthologs (prph2a and 2b) share more than 50% amino acid sequence identity with rom1 orthologs (rom1a and 1b). To test this, we transfected HEK293T cells to express either prph2a-V5 or rom1a-V5 and performed IHC with the prph2 antibody. In support of our hypothesis, both prph2a-V5 and rom1a-V5 showed overlapping signals from the anti-V5 and prph2 antibodies (Fig S3). Thus, the prph2 antibody recognizes both prph2a/b and rom1a/b. As such, we conclude that our double prph2a/2b mutant does in fact lack prph2 and that the remaining signal we detect in rods represents rom1a and rom1b. We therefore refer to this antibody as prph2/rom1 from this point forth.

To fully characterize the phenotype of the double mutant we collected retina samples at 6 timepoints encompassing early development (5 and 15dpf), juveniles (30dpf), as well as adults (90, 180 and 360dpf). Imaging was restricted to sections collected from our standardized anatomical location of the central retina. With the known scarcity of studies observing prph2 loss of function effects on cone PRCs, we first sought to first investigate the consequences of prph2 loss in cones. When observing cone OSs at 5dpf using PNA staining, UV or blue opsin antibodies we confirm the presence of cone OSs, however they appeared to be rounded and stunted (Fig 2C). The average PNA+ cone OS length of the double mutants was just 1.34μm compared to 4.35μm in controls (Fig 2D). The average number of PNA+ cone cells was not affected (Fig 2E). Using TEM to further investigate the ultrastructure of double mutant cone OSs we found the presence of whorl structures which phenocopy cone OS whorls observed in PRHP2^+/-^ mice ^32^ (Fig 2F). All cones observed exhibited the whorl phenotype. By 15dpf we observed very few, if any, cone PRCs compared to controls, which suggests that the whorl structures lead to cone cell degeneration (Fig 2G). By the juvenile stage of 30dpf, we observe only cone OS debris, (Fig 2H). At 3, 6 and 12 months we continue to observe the absence of PNA+ OSs and only what appears like OS debris. We therefore conclude that zebrafish cone cells are critically sensitive to the presence of prph2a and 2b, as they fail to form proper and organized outer segments and undergo degeneration soon after their development in the absence of prph2 function. These phenotypes are in line with what has been observed in studies of murine PRPH2.

### Loss of prph2a and prph2b function triggers futile cycles of cone cell regeneration

When characterizing the effects of prph2a/b loss on cone cells we noted a significant reduction in their number by 15 and 30dpf, but absent of a total reduction (Fig 2G). We hypothesized that this is a result of continuous attempts at regeneration of cone cells. To test this, we used PCNA to measure the amount of dividing and therefore potentially regenerating retinal cells. At 5dpf, we did not observe significant a significant increase in cell proliferation, which is consistent with most cone cells persisting at this stage (Fig S4A). By 15dpf, we saw a significant increase in the PCNA+ cells in the double mutant retina with many of the PCNA positive cells localizing to the ONL (Fig S4A-B). At 30dpf, we continued to observe an enlarged proliferative cell population in the double mutant compared to WT. Collectively, we see a clear trend in the rise of proliferating cells in the double mutant retina in response to degeneration of cone PRCs. This rise of PCNA+ cells in the double mutant retina highlights attempts by the retina to regenerate those lost cone cells which arise, but again degenerate upon failing to assemble proper OS due to the absence of prph2a/b.

### Loss of prph2a and prph2b leads to rod OS enlargement

Having observed consequences of prph2a/b loss of function in cones, we next examined rods. To visualize rod OSs we used our prph2/rom1 antibody paired with wheat germ agglutinin (WGA) In prph2a/2b double mutants the prph2/rom1 antibody staining was consistently observed throughout rod OSs at all timepoints. As mentioned previously in our double mutants this signal represents rom1a/b expression and localization. Prph2/rom1 signal combined with 1d1 staining (rhodopsin antibody at 5 and 15dpf) and WGA (30dpf+) was then used to quantify rod OS length and rod cell number (Fig 3C, D). At 5 dpf, we did not observe any difference in rod OS length or rod cell number between double mutants and controls (Fig 3A-D). Strikingly, by 30dpf not only were the double mutant rod OS present, but they also measured at 45.29μm compared to 17.53μm in WT, an astounding increase of more than 250% (Fig 3B, D). Despite the enlargement there was no difference in the overall number or rod cells between the double mutants and WTs (Fig 3C). This significant overgrowth appears to climax at 30dpf as in latter timepoints, 3mpf to 1ypf, the double mutant rod OS continues to measure between 43-46μm compared to 28-30μm in WT (Fig 3B). Visually, the double mutant rod OSs also appear to encompass the entire OS space compared to the orderly organization of WT retina ONL. (Fig 3A). To determine whether rod OS overgrowth interferes with OS structure we analyzed the ultrastructure of double mutant rod OS using TEM at 30dpf. Results from TEM confirm that rod OSs appear elongated while normal disc curvature is observed (Fig 3E). This suggests that despite the increase in OS length, rod OS appear stable and do not degenerate in the absence of prph2a/b function. Overgrowth of the rod OSs may result from the absence of cone cells with rods filling in the vacant space. This phenomenon has previously been observed in pde6c and prom1b mutant zebrafish ^33,34^.

**Figure 3:**
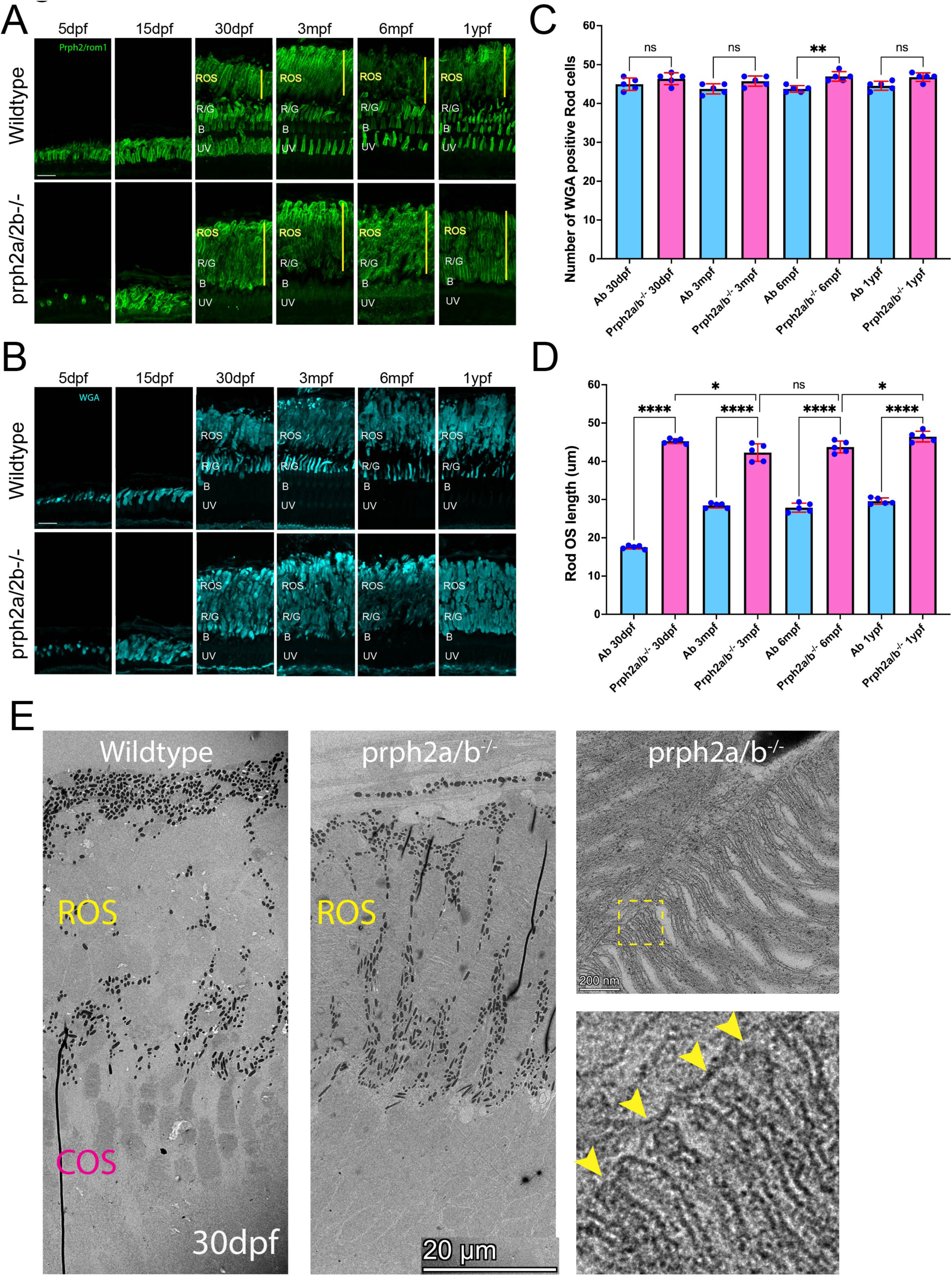
Loss of prph2a and prph2b leads to rod OS overgrowth. **A)** Confocal imaging of central retina cryosections from 5, 15, 30, 90, 180 and 360dpf wildtype or prph2a/b^-/-^ double mutant individuals, probed with anti-prph2 antibody (green). Scale bar = 5μm. ROS = rod outer segments, R/G = red/green cones, B = blue cones, UV = UV cones. **B)** Confocal images of central retina cryosections from 5, 15, 30, 90, 180 and 360dpf wildtype or prph2a/b^-/-^ double mutant individuals, probed with WGA (teal). Scale bar = 5μm. **C)** Quantification of rod cell numbers at 5, 15, 30, 90, 180 and 360dpf comparing wildtype and prph2a/b^-/-^ double mutants. * p<0.05, ** p<0.01, *** p<0.001, **** p<0.0001. **D)** Quantification of rod OS length at 5, 15, 30, 90, 180 and 360dpf comparing wildtype and prph2a/b^-/-^ double mutants. * p<0.05, ** p<0.01, *** p<0.001, **** p<0.0001. **E)** Transmission electron microscopy images of wildtype and prph2a/b-/- retinal sections at 30 dpf. Inset depicts normally formed rims of rod OS discs, highlighted by yellow arrows, in prph2a/b^-/-^ mutants.

To ensure that our prph2b^-/-^ mutant allele was neither hypomorphic or neomorphic, we also generated a second prph2a/2b double mutant line by targeting for deletion the prph2b translation start site in the original double mutant. This line, prph2a/2b^-/-V2^, exhibited nonsense mediated mRNA decay for both prph2a and prph2b as observed by WISH (Fig S5). Comparison of the double mutant V2 line to our original prph2a/2b^-/-^ line confirmed that the prph2b^Δ18–157^ allele is also null as there were no differences in phenotypes observed (Fig S5).

### rom1a and rom1b are necessary to assemble and maintain rod OSs

Based on results from our prph2a/b double mutant line we conclude that in zebrafish prph2a/b function is critical for cone OS formation and maintenance. Conversely, rod OSs remain stable and due to the absence of cone cells increase their OS length (Fig 3). Mechanistically we predicted that rod OS remained stable due to the presence of the prph2’s partners rom1a and rom1b which in zebrafish are exclusive to rod cells (Fig S6A,B). To test this hypothesis, we designed crRNA against rom1a and rom1b and injected them with Cas9 into wildtype and our prph2a/b double mutant embryos (Fig S6C). Injected prph2a/b mutants and wildtype controls were reared until 5 and 10 dpf and analyzed for cone and rod OS morphology. When comparing 5 and 10dpf retinas to un-injected double mutants it was strikingly clear that targeting rom1a and 1b was sufficient to eliminate essentially all prph2/rom1 signal in rod OSs (Fig 4A). Additionally, both wildtype and prph2a/b^-/-^ embryos injected with rom1a/b crRNA/Cas9 exhibited rounded and generally mis formed ROSs (Fig 4A). The shape of ROSs appeared to phenocopy the rounded cone OS observed in prph2a/b^-/-^ double mutants (Fig 2C).

**Figure 4.**
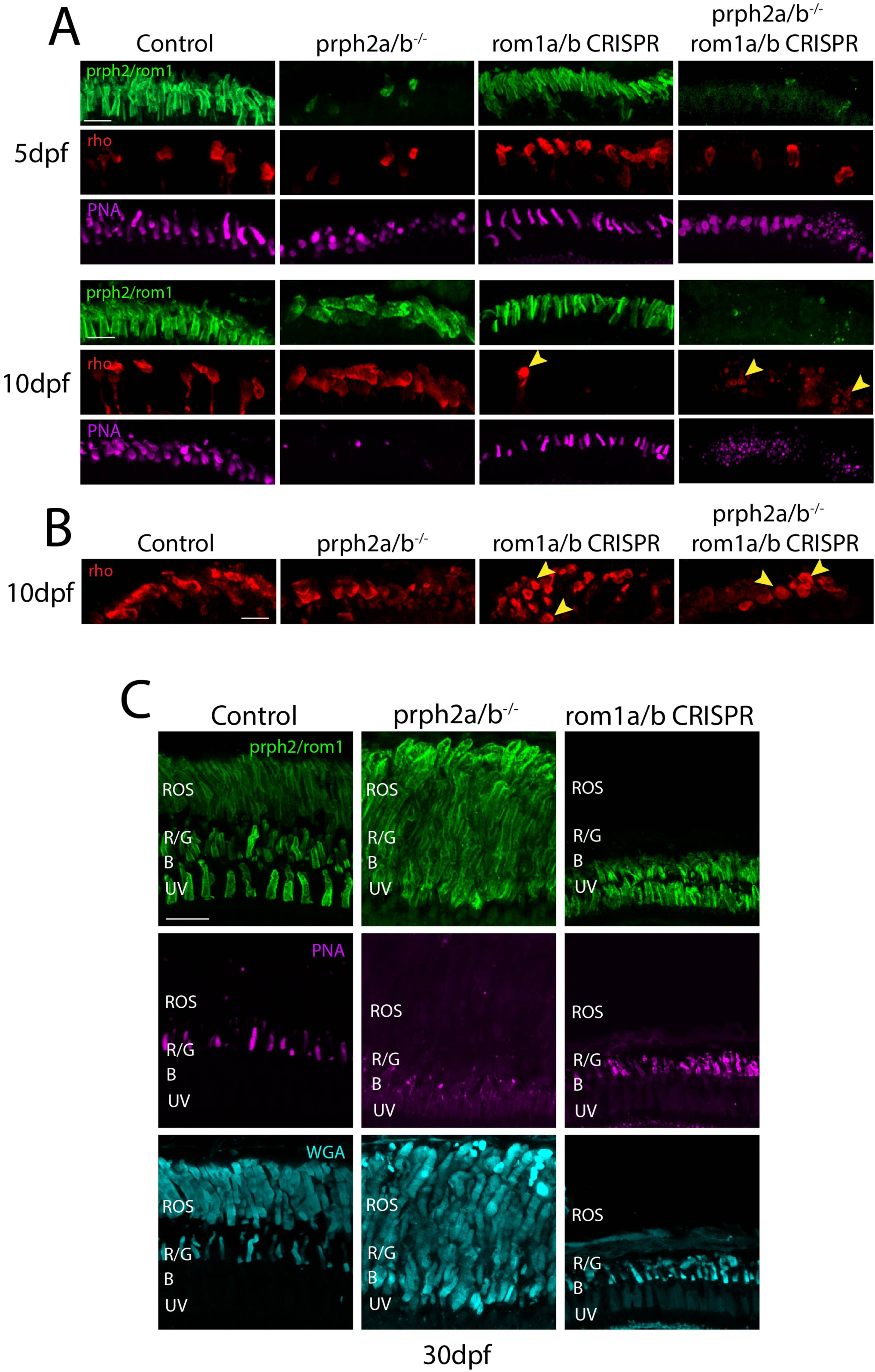
: Inhibition of rom1a/b leads to loss of rod outer segments. **A)** Confocal images of central retina cryosections from 5 and 10 dpf wildtype, prph2a/b^-/-^, rom1a/b CRISPR injected or prph2a/b^-/-^ + rom1a/b CRISPR injected larva probed with anti-prph2/rom1 antibody (green) and anti-rho (1D1) antibody (red). Yellow arrows highlight rounded ROS. Scale bar = 5μm. **B)** Confocal images of peripheral retina cryosections from 10 dpf wildtype, prph2a/b^-/-^, rom1a/b CRISPR injected or prph2a/b^-/-^ + rom1a/b CRISPR injected larva probed anti-rho (1D1) antibody (red). Yellow arrows highlight rounded ROS. Scale bar = 5μm. **C)** Confocal images of central retina cryosection from 30 dpf wildtype, prph2a/b^-/-^ and rom1a/b CRISPR injected zebrafish probed with anti-prph2/rom1 antibody (green), PNA to label red cones (magenta) and WGA to label rod outer segments and green cones (teal). ROS = rod outer segments, R/G = red/green cones, B = blue cones, UV = UV cones. Scale bar = 50μm.

Rounded ROSs were particularly evident near the periphery of the retina where newly formed rod cells are concentrated (Fig 4B). At the same time rom1a/b crRNA/Cas9 injection had no significant effects on cone OSs as assessed by PNA staining (Fig 4A). By 30dpf we continued to observe no effects of rom1a/b crRNA/Cas9 injection on cone OSs yet a near complete loss of all rod OS except those newly differentiated near the ciliary marginal zone which display the rounded OS phenotypes (Fig 4C). To confirm the efficiency of our rom1a/b crRNAs we performed amplicon sequencing of the crRNA target region and detected over 90% of sequences representing indel mutations, thus validating that rom1a and rom1b were targeted and likely inactivated. Taken together, these results suggest that in zebrafish rom1a and rom1b are necessary for assembly of the rod OS while prph2a and prph2b, although present in rods are only necessary for cone OSs

## DISCUSSION

To our knowledge our work here describes the first use zebrafish to model Prph2 loss of function. We show that loss of either prph2a or prph2b does not lead to significant impacts on rod or cone OSs, likely do to ortholog compensation. However, combining the two mutations leads cone OSs with whorls-like structure akin to the OS whorls seen in heterozygous PRPH2 mice ^24,35^ yet with no impact on rod OS assembly. Although the whorl-like cone OSs appear to degenerate by 15 days and are cleared soon after, their presence and the persistence of rod OSs are both in stark contrast to the complete absence of any OS discs in PRPH2 null mice ^24^. The observed cone OS whorl phenotype could be explained by restricted localization of prph2 to cone OS edges adjacent to the connecting cilium (CC) ^18^, the absence of which could lead to impairment of rim curvature at the CC edge thus enabling the discs to grow uncontrollably and forming whorls. Interestingly, PROM1 a cholesterol binding pentaspanin protein has been shown to juxtapose the localization of Prph2 in xenopus cone OS discs and mice nascent rod OS discs ^17,18^. Mutations in PROM1 lead to cone rod dystrophy while complete loss of PROM1 in mice and xenopus PRCs also leads to overgrown OS discs in the shape of OS whorls like those seen in our zebrafish prph2 double mutant ^36–38^. Molecularly, PROM1 consists of five trans membrane domains and has been shown to cause membrane tubulation and protrusions ^39,40^. As such, it is plausible that in cones both PROM1 and PRPH2 work in concert to facilitate proper disk curvature and prevent whorl-like overgrowth.

The other striking result from our prph2a/b double mutant zebrafish was the persistence and significant overgrowth of rod OSs. This is again in stark contrast to the mouse PRPH2 mutants which completely lack rod OS discs and instead only generate OS vesicles^41^. Since ROM1 is a homolog and an interacting partner of PRPH2 with similar functional domains, we predicted the persistence of rod OS in the prph2a/b double mutant may result from functional compensation via zebrafish homologs rom1a and rom1b. In support of this prediction, we show that targeting rom1a/1b using CRISPR/cas9 leads to shortened/stunted rod OSs, which also appeared whorl-like in early development before degenerating. These results confirmed our prediction that rom1a and rom1b can functionally compensate for the loss of prph2a/b in rod PRCs.

Surprisingly, wildtype embryos injected with same rom1a/1b CRISPR/cas9 combination also exhibited significant impairment and loss of rod OSs but not cone OSs which were unaffected. This result was unexpected since ROM1 KO mice do not exhibit severe PRC degeneration compared to the PRPH2/RDS mutants ^42^ and since prph2a/b were not affected (Fig 4). As such, our results indicate that in zebrafish prph2a/b function is critical in cone PRCs but dispensable in rods while rom1a/b is critical for proper rod OS formation instead of prph2a/b. This striking difference in phenotypes between our zebrafish prph2a/b and rom1a/b mutants compared to mice suggests divergent roles of prph2 and rom1 in zebrafish. Taken together this data indicates that in zebrafish cone cells rely solely on prph2a/b function, while rod cells despite expressing both prph2a/b and rom1a/b rely solely on rom1a/b function (Fig 5).

**Figure 5:**
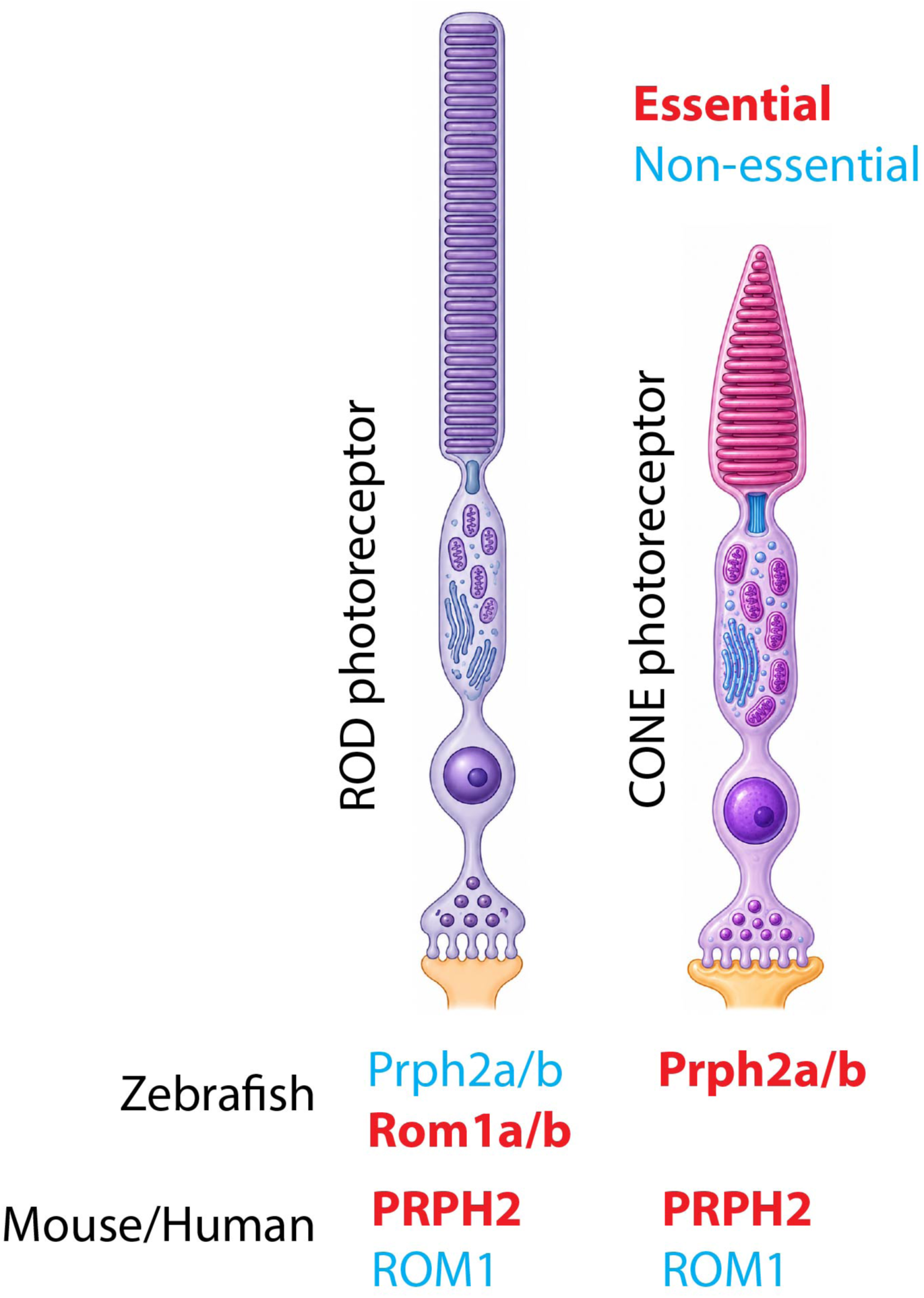
Model for the roles of prph2 and rom1 in zebrafish OS assembly.

The results of our study put a new twist in the evolutionary story of prph2 and rom1 function during photoreceptor outer segment assembly. Studies of mammalian PRPH2 and ROM1 clearly indicate that when it comes to formation of OS discs, PRPH2 is king. In both mouse models and human PRPH2 patients, there is clear correlation between the loss of PRPH2 function and retinal dystrophy. In fact, just the loss of one copy of mouse PRHP2 can lead to highly mis-organized OS in the form of whorls, while a complete knockout results in total failure of photoreceptor OS assembly with no disc formation at all ^35,41^. Additionally, in both mice and humans the loss of ROM1 function appears to have only minor effects on OS formation despite the biochemical association of ROM1 and PRHP2 heterotetramers and oligomers observed during disk rim curvature assembly. One would could therefore extrapolate that the functional relationship between PRHP2 and ROM1 would have remained conserved throughout evolution. The results of our study however challenge that notion. Interestingly, amino acid sequence comparison indicates that zebrafish prph2a/b and rom1a/b share ∼50-54% identity while human and mouse PRPH2 and ROM1 share only 34 and 36% identity respectively. This suggests that during the evolution from fish to mammals rom1 lost its necessity in rod cells while prph2 gained the ability to drive outer segment disc assembly in both rods and cones. Based on these unexpected results, one could postulate that during evolution zebrafish diverged the function of rom1a/b and prph2a/b in rods while losing expression of rom1a/b completely form cones. Alternatively, in mammals, evolutionary pressure could have led to the expression of ROM1 in both cell types but having selected PRPH2 as the critical rim curvature mediator with ROM1 playing a supporting or regulatory yet non-essential role. Interestingly, it has been previously shown that ROM1 can modify the phenotype of PRPH2 mutants. In particular, the absence of ROM1 in combination with PRPH-Y resulted in a rod RP like phenotype rather than a cone phenotype ^43^. Thus, mammalian rod OSs can be sensitive to the absence of ROM1 but not as sensitive as in zebrafish. Considering our findings, future work will focus on the biochemical aspects of zebrafish prph2a/b and rom1a/b.

Do prph2a/b and rom1a/b heterodimerize like they do in mammals and if so with what type of stoichiometry does this differ between cones and rods? Alternatively, how would mammalian PRPH2 or ROM1 behave in a zebrafish rod or cone cell, would they retain their mammalian hierarchy of function or adopt the zebrafish paradigm. Overall, our findings suggest that we may learn more about the function of PRPH2 and ROM1 by studying their homologs in lower vertebrates.

## MATERIALS AND METHODS

### Zebrafish Husbandry and Embryo maintenance

Zebrafish husbandry used in all procedures were approved by the University of Kentucky Biosafety office as well as IACUC Policies, Procedures, and Guidelines (IACUC protocol 2021– 3781). The AB strain was used as wildtype. Embryos were kept at 28°C in E3 embryo media. At indicated times in the study, embryos and adults were anesthetized in tricaine before harvesting the eyes and fixed with 4% PFA in PBS overnight at 4°C.

### Statistical analysis

All the data sets were analyzed using Prizm 10. Data shown on graphs represents individual measurements +/- standard deviation. Each of the data points had an n of 5+ individuals. One-way ANOVA with Tuckey post hoc multiple comparisons analysis was used for assessing direct comparison significance between ages or genotypes. All the graphs depict the mean value.

### Availability of data and materials

All data generated or analyzed during this study are included in this published article. All materials, zebrafish lines, and reagents will be shared upon request after publication.

### Whole mount In Situ Hybridization

WISH was performed as previously described ^44,45^. RNA probes were generated via PCR amplification from 3 dpf cDNA fused to T7 promoter sequence and subsequently transcribed (DIG or FITC labeled) using T7 polymerase (Roche). Primer sequences can be found in table S1. Approximately 1000bp was amplified from the 3’ end of each cDNA. Images were captured using a Nikon Digital sight DS-U3 camera and Elements software. Image adjustment was performed using Adobe Photoshop.

### Cryosection and Immunofluorescence/IHC

Embryos and adult eyes were fixed in 4% paraformaldehyde then washed-out with PBS+ 0.1%Tween-20. Next, the specimens were washed overnight in 10% then 30% sucrose overnight at 4°C. Samples were mounted in OTC and frozen at -80^0^C. Transverse, 10 nm sections were collected, beginning just anterior to and ending posterior to the eye. For imaging and cell quantification, only the sections containing the lens were used for consistency. Immunofluorescence was performed as described previously with addition of a Tris-EDTA based antigen retrieval step prior to serum blocking the specimen ^46^. Post blocking, following antibodies and lectins were used: anti-prph2 (Peripherin – 2, rabbit, 1:100, 18109-1-AP Protein-tech, Rosemont, IL, United States), anti-actin (beta Actin, mouse, 1:100, MA1-140 Thermo Fisher, Waltham, MA, United States) anti-gnb1 (GNB1, rabbit, 1:100, PA5-30046 Thermo Fisher, Waltham, MA, United States), anti PCNA (PCNA, mouse 1:100, MA5-11358 Thermo Fisher, Waltham, MA, United States), anti UV opsin (UV opsin, rat, 1:100 a gift from Dr. Ted Allison (University of Alberta)), anti-Blue opsin (Blue opsin, rabbit, 1:100, DZ41798 Bosterbio, anti1D1 (Zebrafish rhodopsin -1D1 mouse, 1:50 a gift from Dr. James Fadool (Florida State University)). Alexa fluor 488, 555, and 647 conjugated secondary antibodies at a concentration of 1:200 (Thermo Fisher, Waltham, MA, United States) were used with DAPI as a counter stain. Lectins: Peanut Germ Agglutinin conjugated with a 488 fluorophore (1mg/ml, 1:100 #29060 Biotium, San Francisco, CA, United States) and Wheat Germ Agglutinin conjugated with a 405 fluorophore (1mg/ml, 1:50, #29028-1 Biotium, San Francisco, CA, United States) were used to label red-cone and green cone + rod outer segments respectively.

### Cloning

Prph2a coding sequence was amplified from 72hpf cDNA with EcoRI and XhoI linkers, cloned into pGEMT-easy and then subcloned into pCDNA3-V5 using EcoRI and XhoI. Rom1a coding sequence was amplified from 72hpf zebrafish cDNA with HindIII and XhoI linkers (Table S1), cloned into pGEMT-easy and then subcloned using HindIII and XhoI into pCDNA3-V5. Whole plasmid sequencing was used to confirm proper ligation.

### Confocal/SIM imaging

Immunofluorescence imaging was performed as described previously ^46,47^. Processed sections were mounted in Vectashield antifade mounting medium (H-1700-10 Vector laboratories, Newark, CA, United States) then imaged on a Nikon C2 confocal microscope under a 60X 1.4 NA oil immersion objective. For consistency all the images were captured from the central retina to assess all the major types of photoreceptors.

Super resolution microscopy was performed on a Nikon A1R confocal microscope equipped with a Structured Illumination Microscopy module and a CMOS sensor under a 100X 1.49 NA oil immersion objective. Quantification of OS length and number was performed using confocal captured images in Image J. Image adjustments were completed in Adobe Photoshop and figures were assembled using Adobe Illustrator.

### Cell Culture

We received the authenticated cells from ATCC (HEK293T: CRL-11268G-1). Cells were tested bi-weekly for mycoplasma using a PCR-based kit from Thermo Fisher (J66117.AMJ). HEK293T cells were grown in DMEM+10% FBS at 37^0^C and seeded on glass coverslips. Transfection was performed using 1ug plasmid DNA+ PEI as described previously (Patel et al 2026). Coverslips were fixed using 4% PFA for 30min, blocked with BSA, incubated with anti-prph2 (1:500) and anti-V5 antibodies (1:500) and subsequently anti-rabbit Alexa488 and anti-mouse Alexa 647 secondary antibodies (1:1000) respectively. DAPI was used as a nuclear stain. Coverslips were mounted onto slides using Vectashield antifade mounting medium and imaged on a Nikon C2 confocal microscope under a 60X 1.4 NA oil immersion objective.

### TEM

TEM sample preparation was slightly modified from (our cdhr1a paper). First, zebrafish larvae and adult specimens were fixed in 2.5% glutaraldehyde and 2% paraformaldehyde in 0.1M phosphate buffer at 4°C overnight on a shaker. Samples were then washed 5 times with 0.1M phosphate buffer (with 0.1% Tween-20) for 5 times - 10 minutes each to wash out the aldehyde fixatives. Next, samples were placed in 1% Osmium tetroxide solution for 1.5 hours in dark at room temperature on a shaker.

Samples were then washed again with 0.1M phosphate buffer with Tween-20 for 5 times – 5 minutes each and subsequently dehydrated with a series of ethanol washes at 50% (10mins), 70% (10 mins), 80% (10 mins), 90% (15 mins), and two 100% ethanol washes (15 mins each). Next, the specimens were infiltrated with the low viscosity formulated Spurr resin through a series of ethanol:resin combinations 25% Spurr:75% ethanol (30 mins), 50% Spurr:50%ethanol (30 mins), 75% Spurr:25% ethanol (1 hour), and with 100% Spurr resin (overnight at 4°C). The following day samples were exchanged with fresh 100% Spurr resin for 3 hours and then resin was polymerized at 65°C. One-micron thin sections were collected via a glass blade from the resin blocks using Leica EM UC7 and were stained with Toluidine blue to assess the location of samples being sectioned. Once satisfied with the location, ultrathin 70nm sections were collected using a diamond knife (Ultra 45° DiATOME, Quakertown, PA, United States) and placed on slotted EM grids (G2010-Ni Electron Microscopy sciences, Hatfield, PA, United States) covered in a layer of 0.5% formvar. Sections were then stained with a non-radioactive heavy metal UranyLess EM stain (#22409 Electron Microscopy sciences) for 30 mins followed by Reynold’s lead citrate stain for 15 mins. Images were then collected using FEI Talos F200X at 200 keV.

### CRISPR editing

To establish the CRISPR/Cas9 based mutant lines, we ordered predesigned and synthesized ALT-R Crispr crRNA (IDTDNA) targeting prph2a (GGTAACGGTTGCTCACCCAC, ACTCCTCCAAGTATGCGAAA), prph2b-V1 (ACTCAATCTGGGTCATATCC, AAGCGAGTCAAGCTGGCCCA), prph2b-V2 (AGTAACGGTTGCTAATCCAC, ATTTTGACCACCCACAATTC). crRNA-tracrRNA duplexes were constructed before injecting with Alt-R Cas9 v.3 enzyme in single cell staged embryos as described previously ^48^. Genotyping was performed by designing PCR primer oligos (Table S1) upstream and downstream of the expected CRISPR/Cas9 based deletion sites and the resulting difference in sequence size was used to genotype using DNA gel electrophoresis. Exact deletion sequences in prph2a and prph2b were confirmed via Sanger sequencing after outcrossing for two filial generations.

### CRISPANT analysis

rom1a (GGAGCGATCACCTTCACTTT, TAATCTATCACAATGGTCGT), rom1b (TGTTATAGGTCTAAGAGTAT, TTCATTCTAAGGTTCTTTCT) crRNAs were co-injected with Alt-R-Cas9 v.3 into single cell embryos. Confirmation of cutting was performed by designing PCR primer oligos (Table 1) upstream and downstream of the expected CRISPR/Cas9 induced deletion sites, amplifying the region and performing nanopore amplicon sequencing (eurofinsgenomics) of the purified amplicons. Rates of insertion/deletions was quantified using CRISPResso2.

## Competing interests

The authors declare that they have no competing or financial interests.

### Primer Sequences

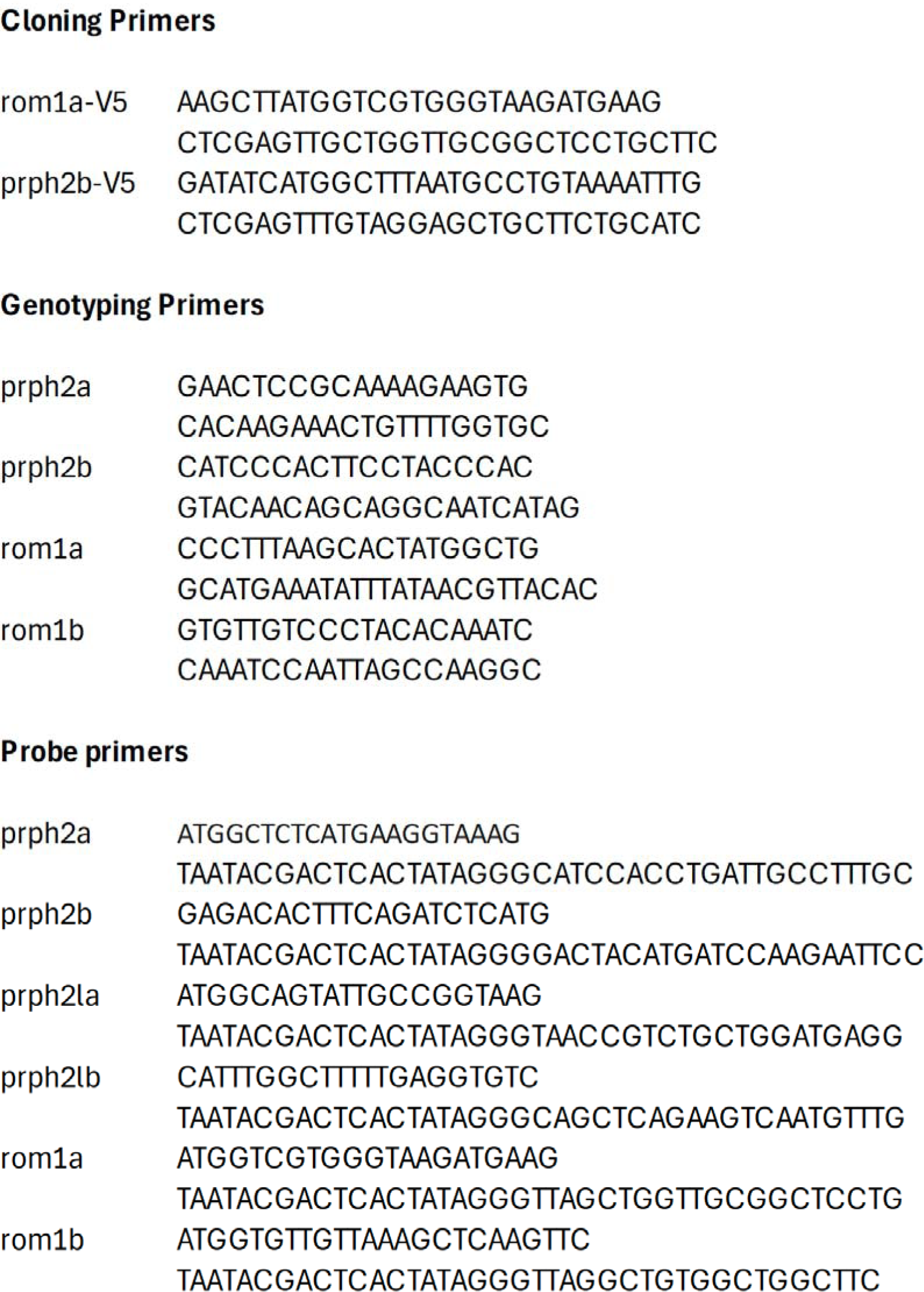

## FIGURE LEGENDS

**Supplementary Figure 1:**
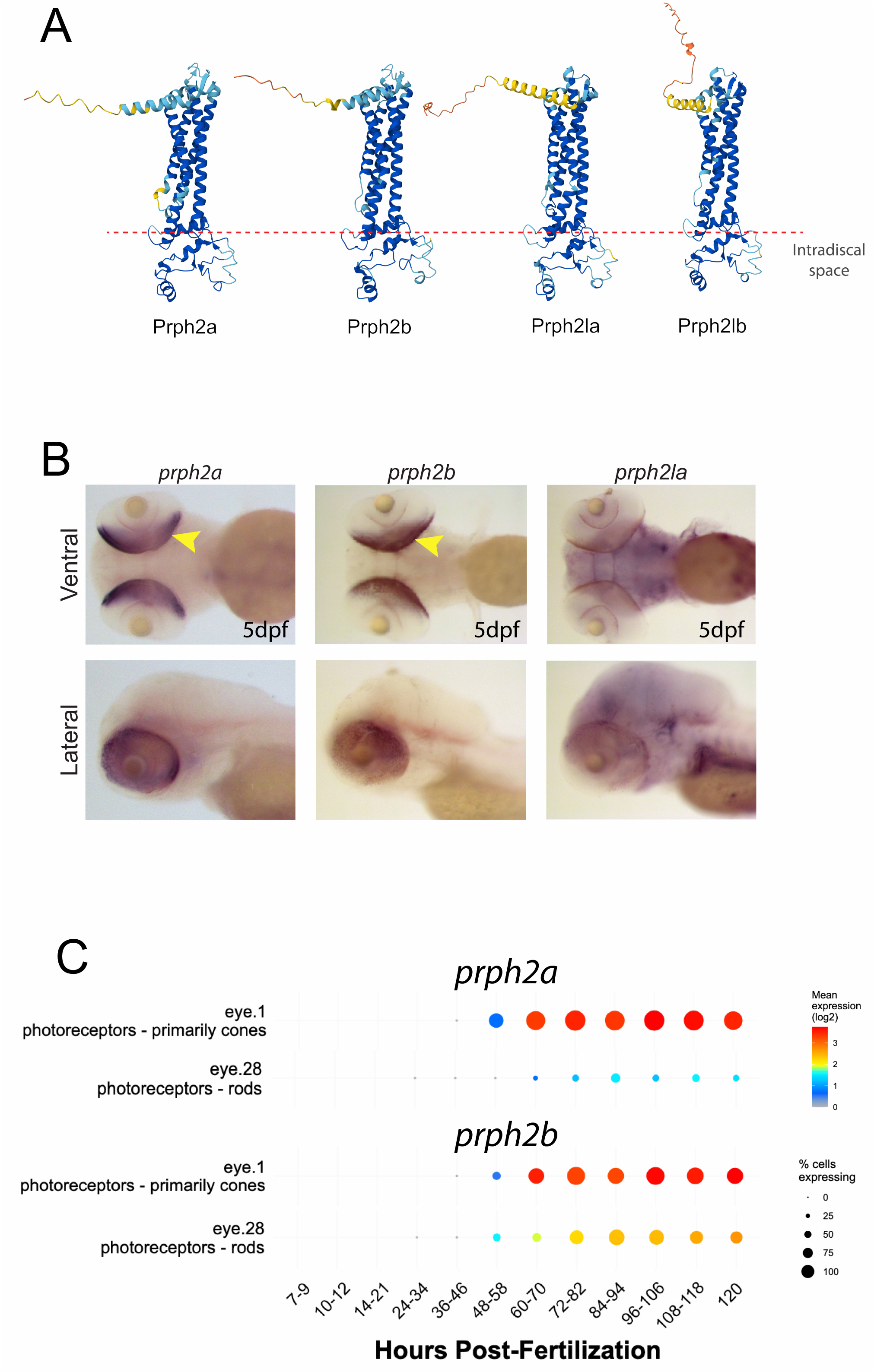
Expression of zebrafish prph2 orthologs. **A)** Alphafold3 models of prph2a, prph2b, prph2la and prph2lb with the intradiscal domain positioned ventrally. Blue color indicates high confidence while red color indicates very low confidence of the model. **B)** Whole mount in situ hybridization of prph2a, prph2b and prph2la probes in 5 dpf zebrafish embryos. Images depict ventral and lateral views of the embryos. Prph2a and prph2b both exhibit expression in the retina (yellow arrows) while prph2la signal is absent from the retina. **C)** prph2a and prph2b developmental expression profiles according to DanioCell. Both genes exhibit robust expression in cones and modest levels in developing rods.

**Supplementary Figure 2:**
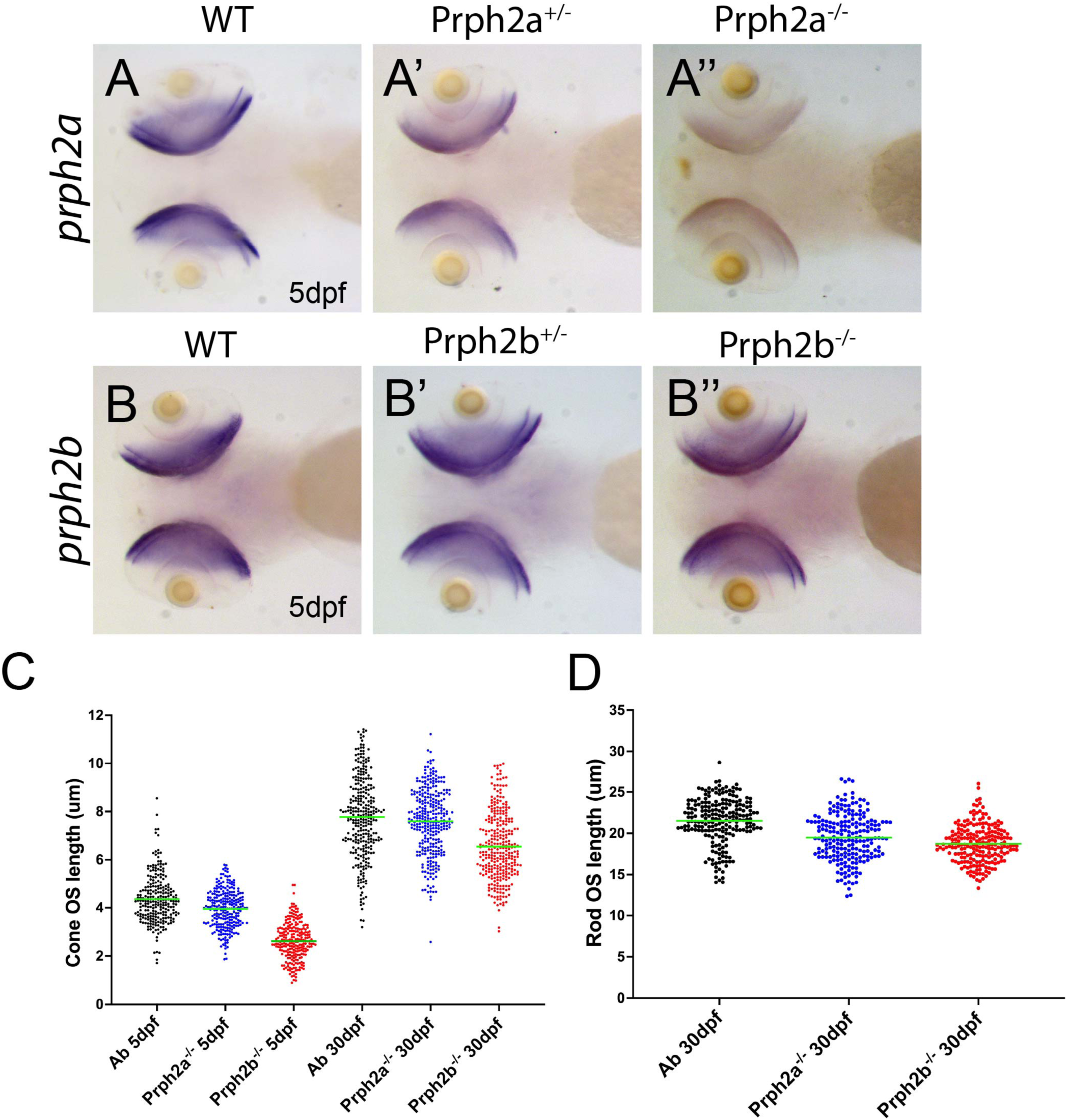
Characterization of prph2a and prph2b mutant lines. **A)** Whole mount in situ hybridization of prph2a probe in 5 dpf wildtype (A), heterozygous (A’) or homozygous (A”) prph2a^-/-^ zebrafish embryos. Images depict ventral and lateral views of the embryos. The prph2a^-/-^ line exhibits non-sense mediated mRNA decay. **B)** Whole mount in situ hybridization of prph2b probe in 5 dpf wildtype (B), heterozygous (B’) or homozygous (B”) prph2b^-/-^ zebrafish embryos. Images depict ventral and lateral views of the embryos. The prph2b^-/-^ line does not exhibit non-sense mediated mRNA decay. **C)** Individual cone OS length measurements of all prph2a^-/-^ or prph2b^-/-^individuals assayed from 5 and 30dpf. **D)** Individual rod OS length measurements of all prph2a^-/-^ or prph2b^-/-^individuals assayed from 30dpf.

**Supplementary Figure 3:**
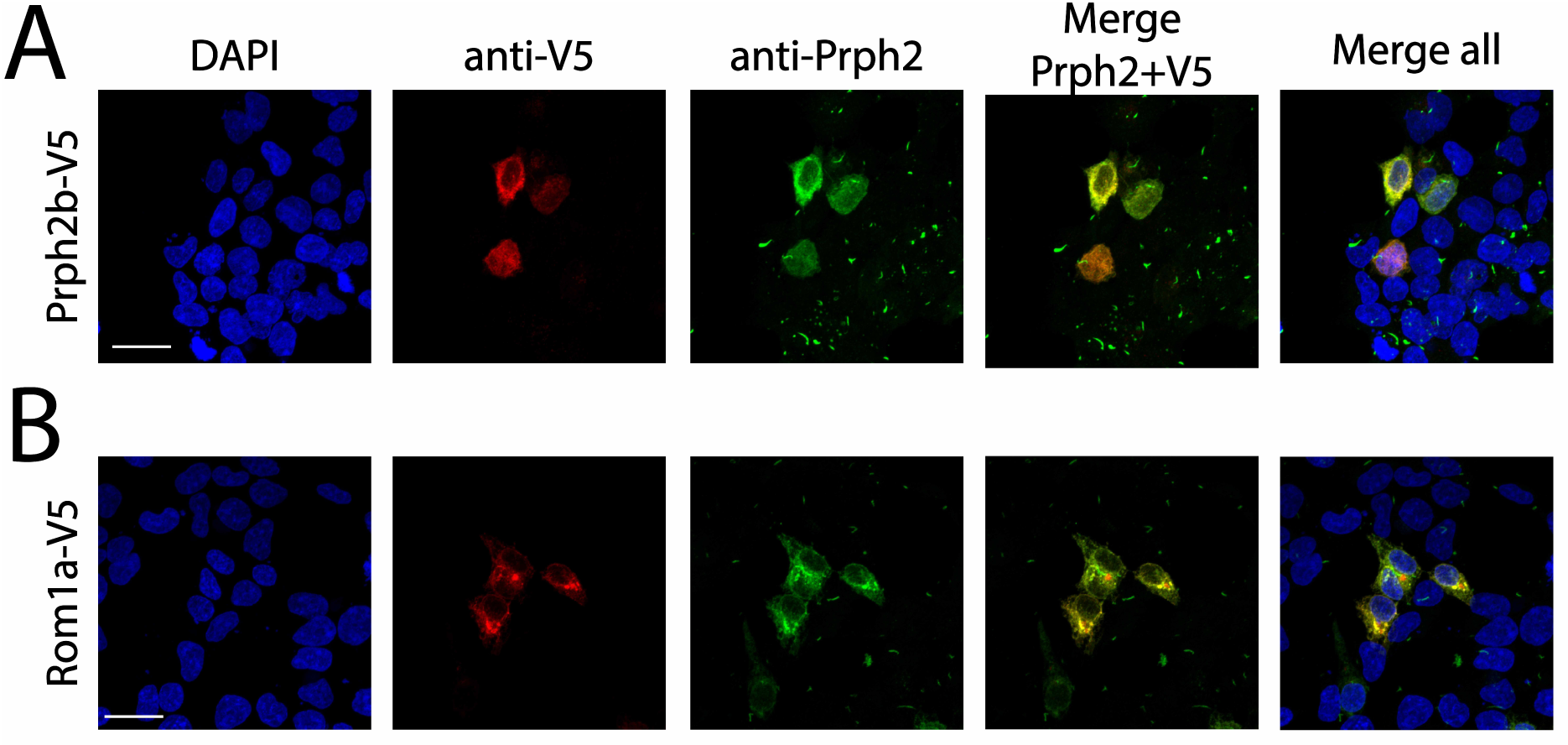
prph2 antibody cross reacts with prph2b and rom1a. **A)** IHC of HEK293 cells transfected with prph2b-V5 probed with anti-V5 antibody (red) anti-prph2 antibody (green) and DAPI (blue). Co-localization of the V5 and prph2 antibody signals is shown in the prph2 + V5 and merge all panels. Scale bar = 10μm **B)** IHC of HEK293 cells transfected with rom1a-V5 probed with anti-V5 antibody (red) anti-prph2 antibody (green) and DAPI (blue). Co-localization of the V5 and prph2 antibody signals is shown in the prph2 + V5 and merge all panels. Scale bar = 10μm

**Supplementary Figure 4:**
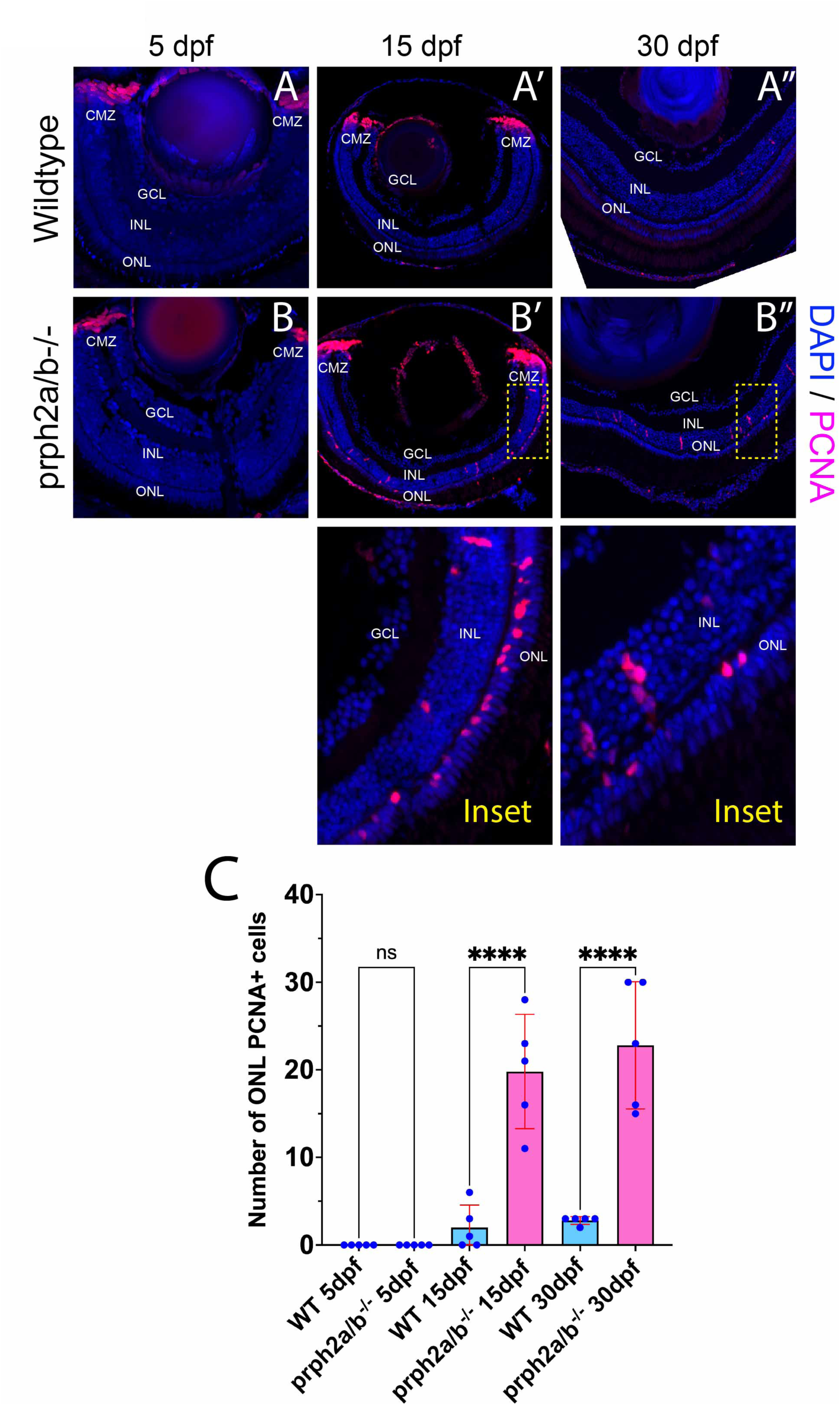
Induction of a regenerative response in the prph2a/b-/-retina. **A)** Confocal imaging of central retina cyrosections from 5, 15 and 30 dpf wildtype individuals probed with anti-PCNA antibody (magenta). ONL= outer nuclear layer, INL= inner nuclear layer, GCL = ganglion cell layer, CMZ= ciliary marginal zone. DNA was stained with DAPI (blue). Scale bar = 100μm **B)** Confocal imaging of central retina cyrosections from 5, 15 and 30 dpf prph2a/b^-/-^ individuals probed with anti-PCNA antibody (magenta). DNA was stained with DAPI (blue). Scale bar = 100μm. Insets depict up-regulation of PCNA in the mutant INL and ONL. C) quantification of PCNA+ cells in the ONL at 5, 15 and 30dpf comparing wildtype to prph2a/b^-/-^ individuals. * p<0.05, ** p<0.01, *** p<0.001, **** p<0.0001.

**Supplementary Figure 5:**
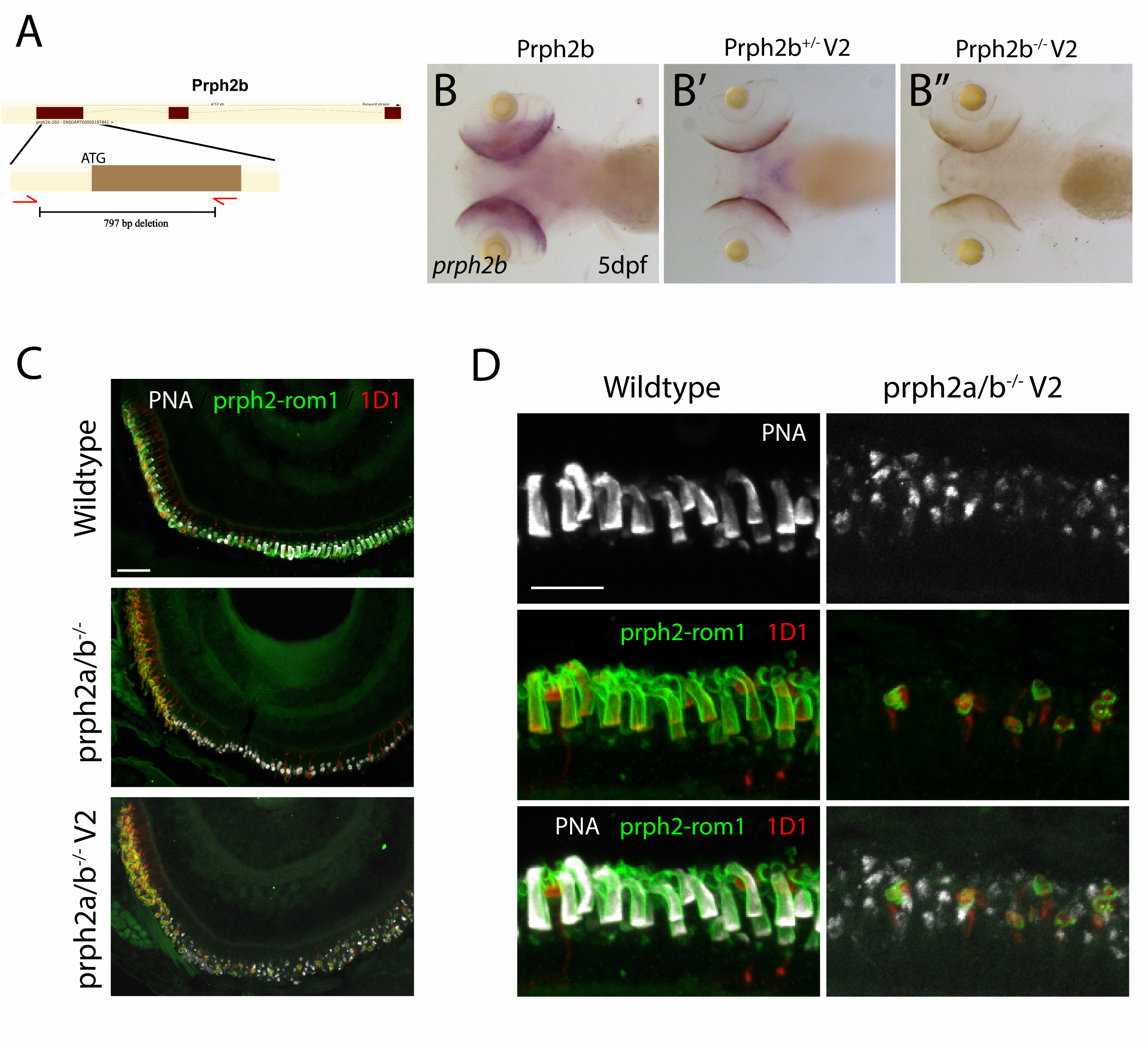
prph2b^v2-/-^ loss of function allele phenocopies prph2b^-/-^. **A)** Schematic diagram of crRNA targeting for prph2b^V2-/-^ allele. **B)** Whole mount in situ hybridization of prph2b probe in 5 dpf wildtype (B), heterozygous (B’) or homozygous (B”) prph2b^v2-/-^ zebrafish embryos. Images depict ventral and lateral views of the embryos. The prph2b^v2-/-^ exhibits non-sense mediated mRNA decay. **C)** Confocal images of central retina cryosections from wildtype, prph2b^-/-^ and prph2b^v2-/-^ 5 dpf individuals probed with PNA (white), anti-rho antibody (red) and anti-prph2 antibody (green). Prph2b^-/-^ and prph2b^v2-/^- exhibit identical phenotypes. Scale bar = 10μm **D)** Confocal images of central retina cryosections at high magnification from wildtype and prph2b^v2-/-^ 5 dpf individuals probed with PNA (white), anti-rho antibody (red) and anti-prph2 antibody (green). Like prph2a^-/-^ the v2 mutant line retains prph2 antibody signal only in remaining rod cells. Scale bar = 5μm

**Supplementary Figure 6:**
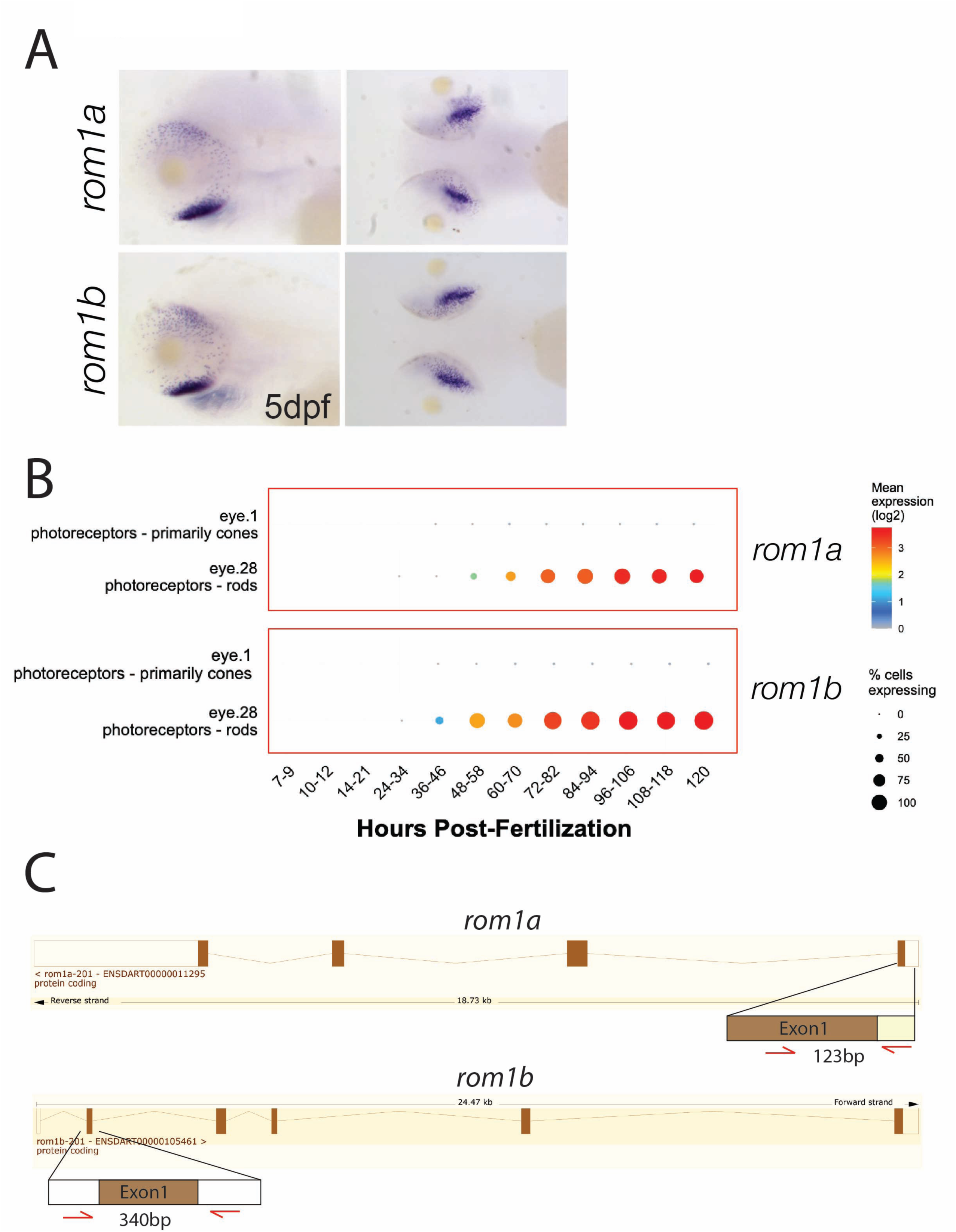
**A)** Whole mount in situ hybridization of *rom1a* and *rom1b* probes in 5 dpf zebrafish embryos. Images depict ventral and lateral views of the embryos. *Rom1a* and *rom1b* both exhibit strong expression in the ventral retina (yellow arrows) reminiscent of rod photoreceptor specific expression patterns. **B)** rom1a and rom1b developmental expression profiles according to DanioCell. Both genes exhibit robust expression in rods and a complete absence of expression in cones. **C)** Schematic outlining the CRISPR-Cas9 targeting strategy for targeting rom1a and rom1b.

